# Multi-level framework to assess social variation in response to ecological and social factors: modeled with coral gobies

**DOI:** 10.1101/2023.11.22.568347

**Authors:** Catheline Y.M. Froehlich, Siobhan J. Heatwole, O. Selma Klanten, Martin L. Hing, Courtney A. Hildebrandt, Jemma O. Smith, Marian Y.L. Wong

## Abstract

Understanding variation in social organization that does not have a strong phylogenetic signal represents a key focus of research in behavioural and evolutionary ecology. In light of this, we established a sociality framework that identifies four categories of variation in social organisation that range from large-scale to fine-scale and can each be related to various ecological factors: (1) forms of sociality, (2) degree of sociality, (3) social plasticity, and (4) within-group plasticity. We modelled this framework by quantifying the four categories of variation over time, space and disturbance regime using multiple species of coral-dwelling gobies from the genus *Gobiodon*. Gobies are a particularly interesting model system as they vary in social structure, have within-group cooperation and form mutualistic relationships with their coral hosts which are vulnerable to climatic disturbances. We found that gobies varied in forms of sociality – from being solitary, to paired or group-living depending on location and disturbance regime. Only low or moderate degrees of sociality were observed in gobies, and this was influenced by location or disturbance regime depending on species. Gobies were more often solitary or pair-forming than group-forming (which became extremely rare) in a high disturbance regime whereas they were more often found in groups in a moderate disturbance regime. The size of coral hosts affected the social plasticity of gobies, and corals were smaller due to climatic disturbances. Gobies did not exhibit within-group social plasticity, as there were no changes to the structure of size-based hierarchies or sex allocation patterns with location or disturbance regime. Lastly, by combining the four categories of variation, we find that there is a high loss of sociality in coral-dwelling gobies due environmental disturbances, which likely affects overall goby survival as living in groups can improve survival and fitness. By using our structured framework, we identified which categories of social variation were influenced by ecological factors like location and disturbance. This framework therefore provides an excellent tool for predicting future responses of animal societies to environmental stressors.

## 2. Introduction

Social living is a common trait in many taxa, with individuals living in groups to gain some type of advantage, such as predation avoidance, improved territory defense, better survival in harsh conditions, increased mate availability, improved habitat quality, and enhanced offspring resilience (Duffy & Macdonald 2010; Firman *et al*. 2020; Hing *et al*. 2017; Nowicki *et al*. 2018; Queller & Strassmann 1998; Rueger *et al*. 2021a). Sociality is often characterized by convergent evolution without a strong phylogenetic signal even between closely related species (Faulkes *et al*. 1997; Hing *et al*. 2019). Instead, group living and social behaviour are often more dependent on ecological pressures that alter the costs and benefits of social living (Duffy & Macdonald 2010; Emlen 1982; He *et al*. 2019; Hing *et al*. 2017).

When considering the impact of social and ecological factors on sociality, it is important to recognize that there are multiple categories of sociality that can be measured at different scales. However, it is often the case that a clear distinction between these categories and scales is not made (Dornhaus *et al*. 2011; Jetz & Rubenstein 2011). For example, a meta-analysis identified that birds in general will live in groups where rainfall patterns are fluctuating at large geographic scales (Jetz & Rubenstein 2011), yet local environmental conditions and smaller taxonomic scales have yielded alternative results (Gonzalez *et al*. 2013). Group size also tends to be the primary measure of sociality in several studies, and yet several species exhibit strong reproductive skew that requires more detailed assessment of the number of breeders and nonbreeders in a group (Avilés & Harwood 2012; Hing *et al*. 2018; Rueger *et al*. 2021a). it is important to take a comparative approach that assesses the effects of both large and small scale factors on the sociality of animal taxa.

Here, we introduce a multi-level sociality framework that identifies four categories of social variation (from large to fine-scale variation) that highlights the extent of sociality amongst a variety of social species. These are (1) forms of sociality (i.e. proportion of individuals that live solitarily, in pairs, or groups), (2) degree of sociality (i.e. whether individuals within a species exhibit one or many forms of sociality ), (3) social plasticity (i.e. ability for group size to shift based on variation in local ecological or social factors), and (4) within-group plasticity (i.e. ability for individuals to change social behaviour and conflict resolution strategies). This framework can be applied to many social taxa and incorporates both ecological and social contexts as predictors for each category of variation (Fig 1). Ecological factors can include both large and small-scale environmental changes. How likely each category of variation will shift in response to predictor variables may affect the survival of individuals, populations or even species as a whole (Booth 1995; East & Hofer 2010; Gil *et al*. 2017; Jordan *et al*. 2009; Strauss & Holekamp 2019). Hence the vulnerability of individuals and populations can be assessed based on how each category of variation responds to predictor factors e.g. whether a taxon will be more social, stay the same, or become less social following environmental challenges. This framework thus provides an outlook of social maintenance, which determines the ability for the taxa to maintain social group living in its entirety, i.e. the structure and the functioning of groups, despite fluctuations in external factors. Thus, elucidating these four categories of social variation will be important for understanding the influences of ecological and social factors on the maintenance of animal societies.

**Fig 1.**
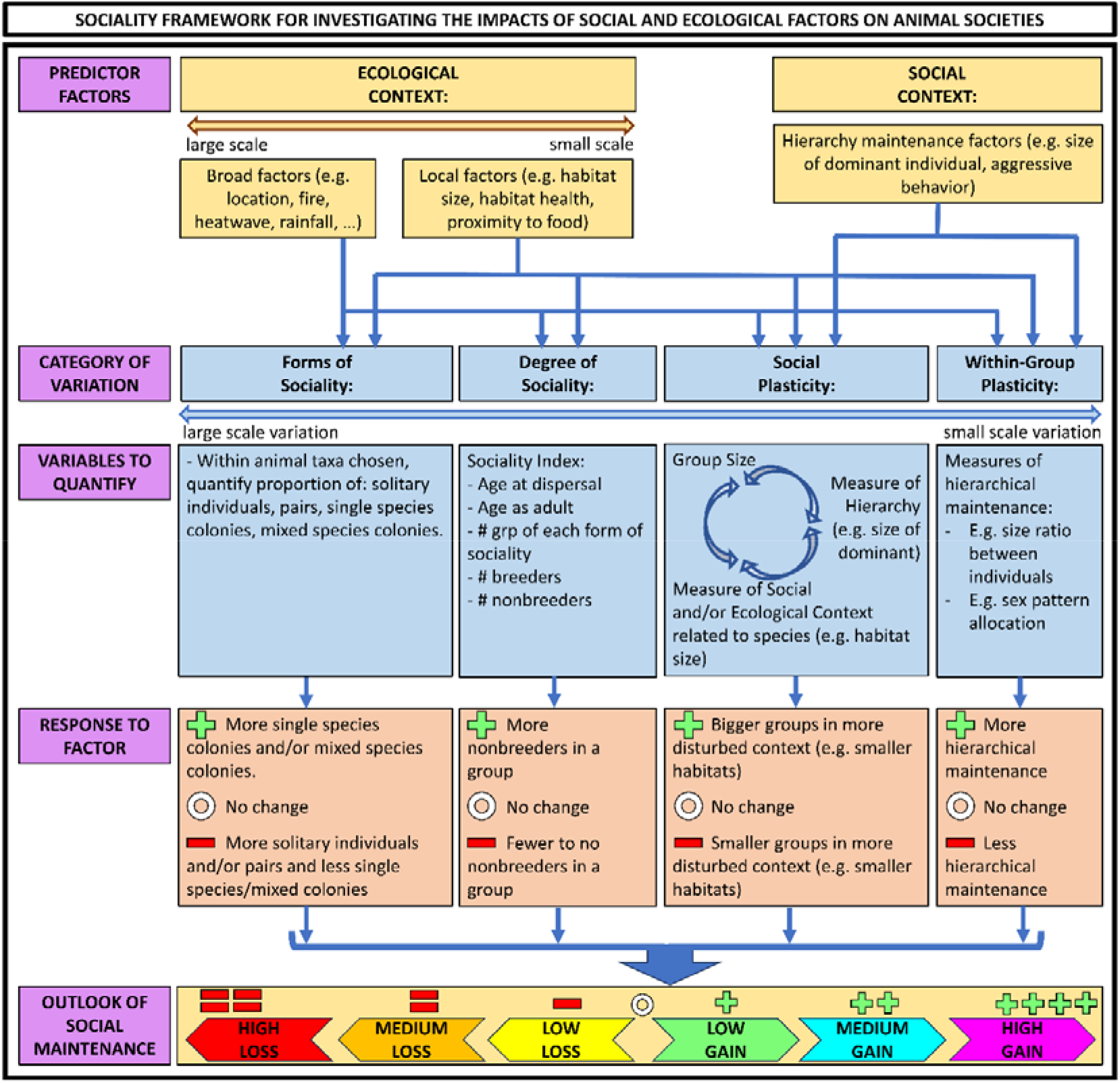
Sociality framework that tests whether ecological factors affect animal societies at four categories of variation and what outlook of social maintenance is given to the taxon based on how many variations have negative or positive responses. Colony = all individual(s) living together in a society; # = number.

At the largest scale, the first category of variation is the form of sociality exhibited, defined as the proportion of individuals in the population that are solitary, in pairs, in single species groups (i.e. >2 group members) or in mixed species colonies (>1 individual of 2+ species of the same taxon) (Fig 1). Note, colony defines any number of individuals (1+) living together, whereas group defines more than 2 individuals living together. The form of sociality can provide an overview of the proportion of individuals living in groups depending on large and small scale factors. The proportion of individuals living solitarily, in pairs, or groups can be affected by ecological conditions, e.g. variability of the environment (Avilés *et al*. 2007; Faulkes *et al*. 1997; Hing *et al*. 2018; Lantz & Karubian 2017) By quantifying the forms of sociality, we can assess whether ecological factors of varying scales will impact the tendency to live solitarily, in pairs or in groups.

The second category of variation is the degree of sociality, defined as the tendency for a species in a given population to be strictly solitary, pair-forming or group-forming. The degree of sociality can be measured via the sociality index, conceived by Avilés and Harwood (2012) using social spiders and mole rats and adapted for fish by Hing et al. (2018). The sociality index provides a value on a scale from 0 (solitary living) to 1 (exclusively group-forming) for a species based on their dispersal, the proportion of groups in a population, and the proportion of breeding and nonbreeding individuals within colonies. Hing et al. (2018) proposed a threshold value of 0.5 to delineate pair-forming and group-forming fish species. The sociality index can be calculated for a species as a whole or for specific populations, depending on what is being tested. Therefore any given species/population is assigned just one value that encompasses how social that species/population is, as well as the extent of reproductive skew exhibited. For a species with the highest degree of sociality, i.e. sociality index close to 1, individuals live strictly in eusocial groups, as seen in naked mole rats, ants, and termites (Avilés & Harwood 2012; Nalepa 2015; Wilson & Hölldobler 2005). Similarly, species with the lowest degree of sociality, i.e. sociality index close to 0, are strictly solitary and hence with low skew, e.g. dune mole rats, platypuses, and solitary sandpipers (Avilés & Harwood 2012; Griffiths 1988; Oring 1973). Values closer to 0.5 are for species that exhibit a mix of social organisation within the population, such as pair-forming and group-forming, e.g. marine shrimp, social spiders, and many birds (Avilés & Harwood 2012; Duffy & Macdonald 2010; Jetz & Rubenstein 2011) with moderate degrees of skew. The degree of sociality therefore provides a value of how social a species is without too much consideration of the extent to which that species could show flexibility in its social arrangements within a set environment, i.e. equivalent to an average degree of sociality exhibited by the species, rather than a variance. The degree of sociality can then be calculated for different populations that vary in ecological conditions (e.g. habitat size, season, latitude/longitude, disturbance regime). Accordingly, the degree of sociality allows us to determine whether a particular form of sociality is consistently the form of sociality that is exhibited by a taxon, or whether ecological conditions allow for different degrees of sociality (Fig 1).

The third and yet finer category of variation in sociality is social plasticity, defined as the extent to which the size of groups within a species or population changes in response to local conditions, such as smaller-scale ecological or social variables (adapted from Teles et al. 2016). For example, within a population, group sizes of some coral-reef fishes vary with the size of their habitat, and in some cases with the size of the largest, most dominant individual within a group (Buston & Cant 2006; Wong, 2011; Rueger et al. 2021). A larger habitat allows more individuals to live together as there is more space and resources, and larger groups in turn can promote an increase in size of the habitat via mutualistically mediated benefits (Buston & Cant 2006; Wong, 2011; Rueger et al. 2021). In addition, the size of the largest individual can dictate the number of smaller subordinates that live within the group owing to rules of the hierarchy (Ang & Manica 2010; Buston & Cant 2006; Wong 2011). Therefore, unlike the degree of sociality which essentially provides just one value to describe overall sociability of a species or population, social plasticity describes how flexible a species or population is to changes in the social and ecological environment, even up to large scale variables like disturbance regimes (Froehlich *et al*. 2021, 2023).

Finally, the finest category of social variation relates to within-group plasticity in sociality, here defined as the extent of conflict and cooperation between individuals within groups and its higher-level consequences through its influence on group structure. In all societies, conflict over rank, resources and reproduction is unavoidable. For some societies, peaceful cooperation by subordinates is maintained through social constraint mechanisms, such as sex, size and maturity regulation, and each of these mechanisms can be influenced by ecological and social factors (Ghiselin 1969; Hing *et al*. 2019; Lassig 1977; Rubenstein 2007; Warner 1988; Wong *et al*. 2008; Wong & Buston 2013). For this category of social variation, the variables that regulate social cooperation can be quantified and related to ecological and social factors. For taxa that exhibit sex allocation patterns, environmental conditions and stressors like rainfall variability, temperature and pollutants have been shown to affect these patterns (Devlin & Nagahama 2002; Oldfield 2005; Ospina-Álvarez & Piferrer 2008; Rubenstein 2007). For example, female superb starlings change their offspring sex allocations based on their own body condition in relation to rainfall variability (Rubenstein 2007). For taxa that exhibit size-based hierarchies, large scale variables like temperature and ocean acidification have been shown to impact some aspects of individual growth (Matthews & Wong 2015; McMahon *et al*. 2019). For example, temperature influences the extent to which subordinates control their own growth in relation to their immediate dominants for Eastern mosquitofish (Matthews & Wong 2015). Small scale variables like habitat size can also affect the growth of individuals depending on their ranks, as seen in hierarchical emerald coral gobies (Wong 2011). Such fine scale variation in social structure can thus be compared among many ecological factors to elucidate whether within-group plasticity exists in relation these factors.

Here, we applied this multi-level sociality framework to understand how and why sociality varies in coral-dwelling gobies from the genus *Gobiodon*, which contains more than 13 species (Munday *et al*. 1999). Within a single colony, defined as all gobies living within a single coral host, gobies have been found living solitary, in pairs, in groups, (Hing *et al*. 2018) and even in mixed species colonies (i.e. with congeners, Froehlich *pers. obs*.) depending on the species. The composition of these mixed species colonies has yet to be quantified, but they provide an additional layer of social complexity as congeners reside and breed within the same habitat and presumably compete for resources. Coral-dwelling gobies likely do not form groups with kin as they have a 3-week larval dispersal stage and then settle into coral colonies as subordinate nonbreeders with unrelated individuals (Brothers *et al*. 1983; Rueger *et al*. 2021b; Wong & Buston 2013). Within groups, individuals are suspected to exhibit peaceful cooperation within a size-based hierarchy, and only a monogamous pair breeds, as seen in the closely related Emerald coral goby *Paragobiodon xanthosoma* (Wong *et al*. 2007). Group sizes mainly depend on ecological factors, like coral size (Hing *et al*. 2019), and potentially on social factors, like body sizes of the largest individual, as seen in *P. xanthosomus* and *Amphiprion percula* (Barbasch *et al*. 2020; Buston 2003; Elliott & Mariscal 2001; Fautin 1992; Rueger *et al*. 2021a; Wong 2011; Wong *et al*. 2007). Within the *Gobiodon* genus, there is only a weak phylogenetic signal for sociality, which suggests that ecological, life history factors may play a substantial role in sociality (Hing *et al*. 2019). *Gobiodon* gobies occur across a range of areas in the Indo-Pacific Ocean, which allows us to test the influences of both large-scale ecological factors, like extreme cyclones and heatwaves, and small-scale factors, like coral size, on the structure of their societies (Froehlich *et al*. 2021; Hughes *et al*. 2018; Munday *et al*. 1999).

Specifically, we investigated how and why sociality varies by examining each of the four categories of social variation in these coral gobies. We used data spanning multiple time points and three different geographic locations which experienced varying disturbance regimes. To use the framework, we (1) compared the forms of sociality exhibited across the *Gobiodon* genus among coral size, time, location and disturbance regimes. We then (2) assessed the impacts of these factors on the three other categories of variation - the degree of sociality, social plasticity, and within-group plasticity - for each individual species and then performed comparisons of these variables among species. Then, we took a closer look at mixed species colonies and investigated which species composed these colonies and quantified the within-group plasticity of these colonies among locations and disturbance regimes. Finally, we combined the results of each sociality metric to identify the outlook of social maintenance of coral-dwelling goby in the face of shifting environmental conditions. (Fig 1).

## 3. Methods

### 3.1. Site Description

The study was conducted at three different locations in the Indo-Pacific, the northern, central and southern locations. The northern location is made up of four inshore sites in Kimbe Bay, West New Britain, Papua New Guinea (PNG) (-5.42896°, 150.09695°). This PNG location has remained relatively undisturbed since an initial trip we conducted in Sep-Nov 2018. The central location is made up of multiple small sites around Lizard Island (LI), Queensland, Australia (-14.687264°, 145.447039°). The LI reef was relatively undisturbed in early 2014 but was affected by four extreme climatic disturbances on an annual basis: category 4 cyclones Ita (2014), Nathan (2015), and two mass bleaching events (2016 and 2017). More recently, LI has sustained mild bleaching events (2020, 2021, and 2022, with only a few patches of corals bleaching) and is in a continued state of disturbances with little time for proper recovery (Froehlich pers. obs., Pratchett et al. 2021). The southern location is within an enclosed lagoon at One Tree Island (OTI), Queensland, Australia (-23.506565°, 152.090954°). The OTI location was relatively undisturbed in 2019 but suffered from mass bleaching events in 2020 with very minimal bleaching in 2022.

### 3.2. Sampling Techniques and Intervals

All fieldwork was conducted either on SCUBA or snorkel at each location. Two types of sampling techniques were used for the study. The first technique involved conducting surveys along 30 m line transects to search all corals within 1 m on either side of the transect. The second sampling technique involved haphazardly sampling corals at each location, and only corals with a minimum of 10cm average diameter were included. When a coral was encountered, a bright torch light (Bigblue AL1200NP) was used to search for goby occupants. Within each coral, the number of gobies (i.e. group size), life stage of gobies, and goby species were noted. Goby life stages were recorded as either breeding adults (two largest adults), nonbreeding adults (all other adults smaller than the two breeders but larger than juveniles), and juveniles (a.k.a. recruits) depending on their coloration and size. Coral diameter was measured along three axes (length, width, and height), and an arithmetic average was taken to indicate coral size (i.e. average coral diameter; Kuwamura et al. 1994). Gobies were collected from a random selection of corals for each sampling technique to quantify body size. During collection, a clove oil anesthetic solution (clove oil, 70% ethanol, and seawater) was sprayed over the coral and fish were wafted out with hand nets (Munday & Wilson 1997). Each fish was placed in a Ziploc bag full of seawater and measured for standard length (mm, ± 0.1 mm) using handheld calipers. During later collections (as noted below), fish were also sexed and injected with a unique visible implant elastomer identification tag (Northwest Marine Technology, Inc., Anacortes, Washington, USA) (Munday 2001). Fish were then returned unharmed to their coral. On later trips, goby colonies containing tagged fish were revisited and re-collected to note coral size, group size, fish size and sex.

Sampling was completed at LI before climatic disturbances (Feb 2014) and three years after the four major climatic events (Jan-Mar 2020). During 2020, gobies were tagged with elastomer and sexed, and then the same colonies were revisited one and two years later (Jan-Mar 2021 and Jan-Apr 2022). Haphazard sampling was completed at PNG during one sampling event (Sep-Nov 2018) in which gobies were tagged with elastomer and were revisited six months later (May-June 2019). Haphazard sampling was completed at OTI before climatic disturbances (Jan-Feb 2019) and two years later (Mar-Apr 2022) after mass coral bleaching had occurred.

### 3.3. Data Analysis

#### 3.3.1. Form of Sociality – Single Species Pairs, Single Species Groups, Mixed Species

Gobies encountered during transect surveys and haphazard searches were included for analysis and were categorized into form of sociality as follows: one individual living alone (solitary), living in pairs with conspecifics only (single species pairs), living in groups with conspecifics only (single species groups), and living with congeners (mixed species). Only corals with a minimum of 10 cm average diameter were included because that was the minimum size of hosted corals measured during haphazard searches. The effect of location (fixed factor) on the form of sociality of gobies were analysed using multinomial logistic regression models for three analyses: (1) compare locations in relatively undisturbed conditions (i.e. before climatic disturbances = PNG2018, LI2014, OTI2019), (2) compare locations before and after being disturbed by climatic disturbances (i.e. pre-disturbances = LI2014 & OTI2019, post-disturbances = LI2020, LI2022 & OTI2022) and the interaction between location and pre/post-disturbances, and (3) compare LI between the two post-disturbance time points (LI2020 and LI2022) to assess recovery. For each multinomial model, the baseline reference level for the response variable was a solitary individual. Juveniles were included in the analysis if they were found with at least 1 adult, as juveniles tend to move between corals if solitary. All analyses were completed in R (v4.2.0) (R Core Team 2022) and R Studio (2022.02.2+485) (RStudio Team 2022) with the following packages: tidyverse (Wickham *et al*. 2019), VGAM (Yee 2010), car (Fox & Weisberg 2019), and rcompanion (Mangiafico 2016).

#### 3.3.2. Degree of Sociality - Sociality Index

We calculated the sociality index for each species in which there were a minimum of 5 colonies of the species in any single location at each survey time point, including pre- and post-disturbance. The sociality index was adapted from Avilés and Harwood (2012) as follows:

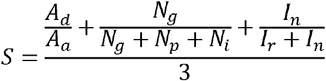

where *A*_*d*_ = age of dispersal, *A*_*a*_ = age of adulthood, *N*_*g*_ = number of groups, *N*_*p*_ = number of pairs, *N*_*i*_ = number of solitary individuals, *I*_*n*_ = number of reproducing (dominant) adults, *I*_*n*_ = number of non-reproducing (subordinate) adults. The numerator is comprised of three components: the proportion of the life cycle spent in a colony, the proportion of groups encountered, and the proportion of subordinates (nonbreeding) individuals (respectively). We followed guidelines set out in Hing et al. (2018) to calculated biologically-relevant assumptions of the numerator. Accordingly, we set the maximum proportion of life cycle spent in a colony (i.e. *A*_*d*_*/A*_*a*_) to 1, which is biologically realistic even if there is some natural variation, as gobies spend only 22-41 days in the larval dispersal stage (Brothers *et al*. 1983). We then calculated the sociality index for each species at each location and time point, and categorized them as either pair-forming (< 0.5) or group-forming (≥ 0.5), as per the threshold of 0.5 (Hing *et al*. 2018). Note, we did not calculate sociality indices for mixed species colonies as colonies were not always made up of the same species combination.

#### 3.3.3. Social Plasticity: Group Size – Size of the Dominant – Coral Size

To investigate the determinants of social plasticity, we only calculated the relationship for goby species that were group-forming as per sociality indices (i.e. >0.5; (Hing *et al*. 2018)), and for which we collected a minimum of 30 colonies. We excluded any mixed species colonies. The analysis of the synergistic relationship between group size, size of the dominant individuals and coral size was repeated for each variable by placing each as the focal response variable in the model. The effect of the size of the dominant individual and coral size on group size were analysed using a generalized linear model using the poisson distribution. The effect of the group size and coral size on the size of the dominant individual were analysed using a linear model. The effect of the size of the dominant and group size on the coral size were analysed using a linear model. Location was included as a fixed factor in each analysis and analyses were repeated separately per species. The variables and models were assessed for normality and homoscedasticity via Q-Q plots, histograms, and residuals over fitted plots, and were transformed as required. If outliers fell outside of 2.5 standard deviation from 0, then they were subsequently removed. All analyses were completed in R (v4.2.0) (R Core Team 2022) and R Studio (2022.02.2+485) (RStudio Team 2022) with the following packages: tidyverse (Wickham *et al*. 2019), lme4 (Bates *et al*. 2015), lmerTest (Kuznetsova *et al*. 2017), LMERConvenienceFunctions (Tremblay & Ransijn 2020), piecewiseSEM (Lefcheck 2016), and emmeans (Lenth *et al*. 2020).

#### 3.3.4. Within-group plasticity: Size Ratios

To investigate within-group plasticity, we investigated the influence of several factors on the size ratios of fish, as size ratios are indicators of peaceful cooperation within size-based hierarchies. For size ratios, we only included single species colonies for which all individuals were collected, otherwise we would not have been able to confirm the correct rank placement of each individual in the hierarchy. Size ratios were calculated by dividing the standard length (SL) of the lower rank (more subordinate) individual by the standard length of the upper rank individual (its immediate bigger group member) (e.g. SL_rank2_ /SL_rank1_) (Wong *et al*. 2007). Size ratios were analysed separately for the breeding pair (i.e. rank 1 and rank 2) as their body sizes were predicted to converge to improve overall reproductive output (Munday *et al*. 2006). The effect of coral size (covariable), group size (covariable), species (fixed factor) and location (fixed factor) on the size ratio between rank 1 and rank 2 individuals (i.e. rankstep 1) were analysed with generalized linear models with family quasibinomial. The analyses were repeated for the size ratio between rank 2 and rank 3 individuals (i.e. rankstep 2) as the next rank after the breeding pair was expected to remain smaller in order to reduce conflict (Wong *et al*. 2007). At two locations, goby colonies were revisited in consecutive sampling events (PNG 2018 & 2019, LI 2020 & 2021); for these repeat visits, size ratios were calculated for rankstep 1 but not for further ranks as there were not enough colonies with minimum of 3 individuals per species. The effect of coral size (covariable), group size (covariable), species (fixed factor), location (fixed factor), and year (fixed factor) on the size ratio for rankstep 1 was analysed with generalized linear models with family quasibinomial.

We had enough samples to compare size ratios of rankstep 1 at LI pre-(2014) and post-disturbances (2020 and 2021). Accordingly, we investigated the effects of coral size (covariable), group size (covariable), species (fixed factor) and pre-vs. post disturbance (fixed factor) on the size ratios of rankstep 1 at LI. The variables and models were assessed for normality and homoscedasticity via Q-Q plots, histograms, and residuals over fitted plots, and were transformed as required. If outliers fell outside of 2.5 standard deviation from 0, then they were subsequently removed. Models were selected based on the Akaike Information Criterion (AIC). All analyses were completed in R (v4.2.0) (R Core Team 2022) and R Studio (2022.02.2+485) (RStudio Team 2022) with the following packages: tidyverse (Wickham *et al*. 2019), lme4 (Bates *et al*. 2015), car (Fox & Weisberg 2019), LMERConvenienceFunctions (Tremblay & Ransijn 2020), piecewiseSEM (Lefcheck 2016), emmeans (Lenth *et al*. 2020), and ggpubr (Kassandra 2020).

#### 3.3.5. Within-group plasticity: Sex Dominance in Breeding Partners

To investigate another aspect of within-group plasticity, we investigated their sex dominance in breeding partners as social coral reef fishes generally have female- or male-dominated societies (Wong & Buston 2013). For single species colonies that were revisited at LI in 2020 and 2021, the sex of the dominant individual (rank 1) was identified on repeated trips. The sex ratio of rank 1 males to rank 1 females was compared to determine whether it differed from 1:1 ratio with a 1-sample proportions test with continuity correction. The effects of species (fixed factor) and year (fixed factor) on the ratio of rank 1 females to rank 1 males within breeding partners was analysed using generalized linear models with the binomial family. All analyses were completed in R (v4.2.0) (R Core Team 2022) and R Studio (2022.02.2+485) (RStudio Team 2022) with the following packages: stats (R Core Team 2022), car (Fox & Weisberg 2019), and rcompanion (Mangiafico 2016).

#### 3.3.6. Mixed Species Colonies: Social Structure and Composition

Across all locations, goby colonies containing mixed species were used to calculated three categorical response variables that measured whether the mixed species colony: (1) had different species intermixed within hierarchical ranks—e.g. rank 1,3,5 were species A and rank 2,4,6,7 were species B (yes, intermixed) versus rank 1-4 were species A and rank 5-7 were species B (no, not intermixed); (2) had the larger-bodied species as the rank 1 individual (yes or no; larger-bodied as defined by Hing et al. 2019); and (3) was composed of solitary individuals, pairs or groups of each species, or a combination of each. The main effect of location (fixed factor) was examined using separate multinomial logistic regression models for each response variable. As mixed species colonies were not collected post-disturbance, body sizes could not be measured and hence no pre-versus post-disturbance analyses were conducted for response variables 1 and 2. When comparing the composition of mixed species colonies (response variable 3) pre-versus post-disturbances, there were insufficient mixed species colonies post-disturbance at LI, hence this analysis is restricted to OTI. The effect of pre-vs. post-disturbance (fixed factor) on mixed species colonies at OTI (response variables 1-3) were analysed using multinomial logistic regression models. For each multinomial model, the baseline reference level for the response variable was as follows: (1) intermixed rank reference: no, (2) larger-bodied species as rank 1 reference: no, and (3) mixed composition reference: solitary individuals. Juveniles were included in the analysis unless they were solitary individuals because juveniles have been seen jumping between different corals when solitary. All analyses were completed in R (v4.2.0) (R Core Team 2022) and R Studio (2022.02.2+485) (RStudio Team 2022) with the following packages: tidyverse (Wickham *et al*. 2019), VGAM (Yee 2010), car (Fox & Weisberg 2019), and rcompanion (Mangiafico 2016).

In addition, we analysed variation in size ratios between adjacent ranked group members. Size ratios for each rankstep within mixed species colonies were calculated up to rankstep 8 due to large group sizes in mixed species colonies. Initially, size ratios were calculated for each species separately within mixed species colonies to test whether size ratios were equivalent to those in single species colonies. The effect of coral size (covariable), group size (covariable), rankstep (fixed factor), species (fixed factor), location (fixed factor) and single vs. mixed species group (fixed factor) on the size ratios (separated by species in mixed species colonies) was analysed with a generalized linear model with family quasibinomial.

Then, size ratios between adjacent ranks were calculated regardless of species, as we confirmed that individuals in mixed species colonies were sometimes intermixed by size within the hierarchy (as determined in the analysis above). The effect of coral size (covariable), group size (covariable), rankstep (fixed factor) and location (fixed factor) on the size ratios (regardless of species in mixed species colonies) was analysed with a generalized linear model with family quasibinomial. Both size ratio models were assessed for normality and homoscedasticity via Q-Q plots, histograms, and residuals over fitted plots, and were transformed as required. If outliers fell outside of 2.5 standard deviation from 0, then they were subsequently removed. All analyses were completed in R (v4.2.0) (R Core Team 2022) and R Studio (2022.02.2+485) (RStudio Team 2022) with the following packages: tidyverse (Wickham *et al*. 2019), lme4 (Bates *et al*. 2015), car (Fox & Weisberg 2019), LMERConvenienceFunctions (Tremblay & Ransijn 2020), piecewiseSEM (Lefcheck 2016), emmeans (Lenth *et al*. 2020), and ggpubr (Kassandra 2020).

## 4. Results

The abundance of *Gobiodon* species differed at each location and some species were found in low abundance at a given location. For example, a latitudinal gradient in opposite directions was previously reported for *Gobiodon histrio* and *Gobiodon erythrospilus* (Munday *et al*. 1999), which we also observed in the current study; i.e. *G. histrio* occurred at PNG and LI (lower latitude) but was extremely rare at OTI (higher latitude), whereas *G. erythrospilus* was never found at PNG but occurred at LI and OTI. Therefore, not all species could be used in each analysis.

### 4.1. Form of Sociality – Single Species Pairs, Single Species Groups, Mixed Species

We compared the form of sociality exhibited by gobies among locations by comparing the proportion of corals that had gobies living alone (i.e. solitary), living in pairs with conspecifics (i.e. single species pairs), living in groups with conspecifics (i.e. single species groups), and living with congeners (i.e. mixed species). We used all species observed for these analyses. Before any climatic disturbances, the form of sociality differed among locations (see Suppl. Tabs 6.1-4 for all statistical outputs, here Suppl. Tab 1, p < 0.01). There were far more mixed species colonies at OTI than any other location, in contrast were more single species groups at LI than at other locations (Fig 2). Beyond these differences, single species pairs were most common at each location (Fig 2). As coral size increased, there was a shift from solitary to single species pairs, to mixed species colonies then finally to single species groups (p < 0.01, Supp Fig 1).

**Fig 2.**
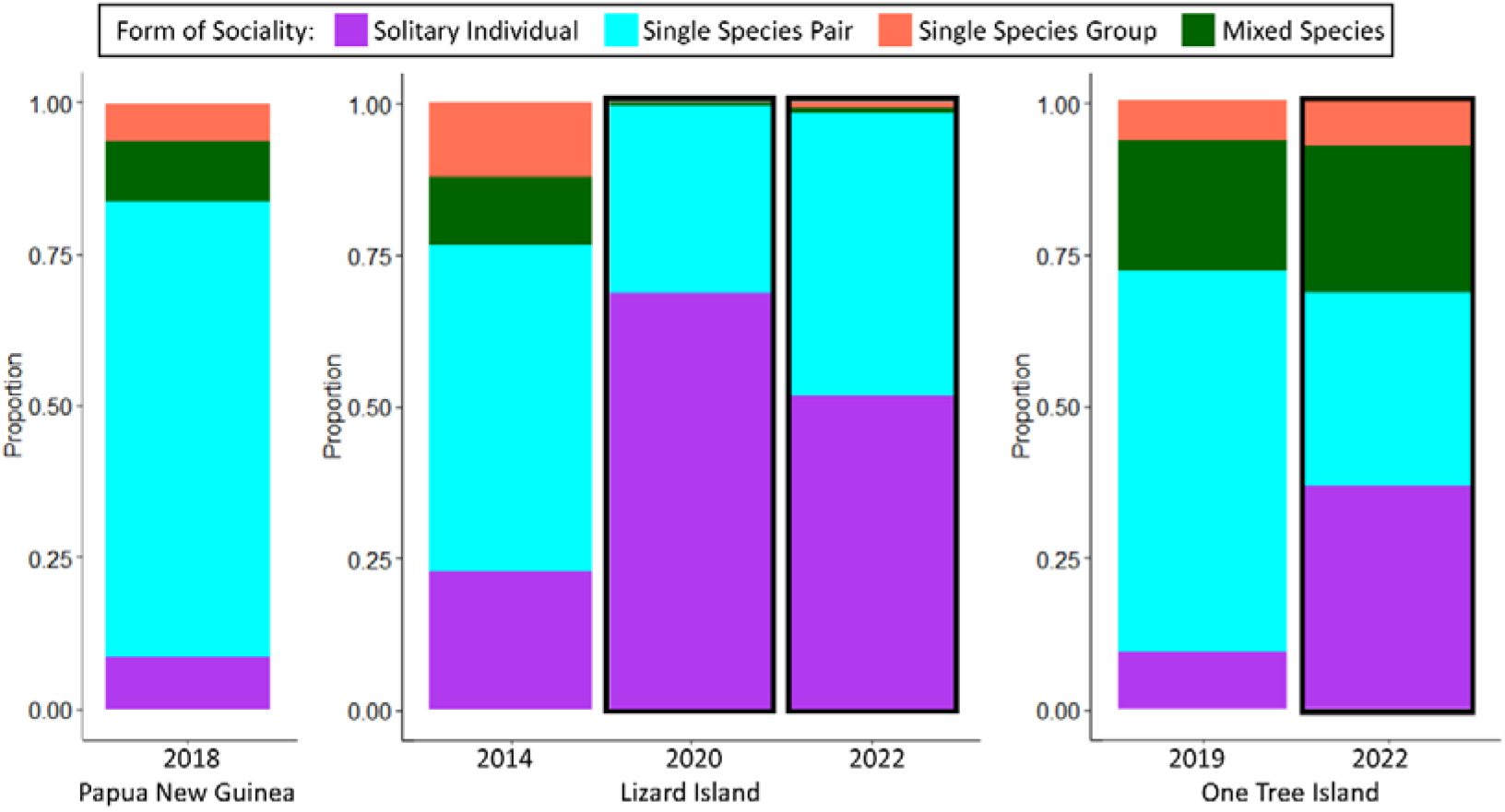
Forms of sociality of all species at all three locations and pre-/post-disturbances for two locations. Data outlined in black line is post-disturbance(s).

There was a significant interaction between location and pre/post-disturbances on the form of sociality (p < 0.01, Fig 2). At OTI, there was a substantially higher proportion of solitary individuals and reduced proportion of pair-forming individuals post disturbance compared to pre-disturbance, but the proportion of single species groups and mixed species colonies remained similar pre- and post-disturbance (Fig 2). At LI, there were also a substantially higher proportion of solitary individuals and reduced proportion of single species pairs post-disturbance than pre-disturbance, but single species groups and mixed species colonies became extremely rare post-disturbance even though that differed slightly among 3-yr and 5-yr mark post-disturbance (p < 0.001, 2020 v. 2022, Fig 2). In 2020, **∼**70% of gobies were solitary compared to just under 25% pre-disturbances, and the remainder were pair-forming except for a single occurrence of a mixed species colony. In 2022, there was a reduced proportion of solitary gobies (**∼**50%), and others lived in pairs except for 5 single species groups (1%) and 5 mixed species colonies (1%). At PNG, there was a similar proportion of solitary and paired individuals as at OTI pre-disturbances, but there was only a slightly higher proportion of mixed species colonies than single-species colonies (Fig 2).

### 4.2. Degree of Sociality - Sociality Index

By calculating sociality indices among locations for each species (minimum of 5 colonies) we found that pair-forming species exhibited low degrees of sociality and remained pair-forming as per Hing *et al*. (2018), even post-disturbances (Fig 3). For species distinctly pair-forming, their index value equaled 0.33 which is the value when a species only ever occurs in pairs. Interestingly, *Gobiodon quinquestrigatus* was defined as pair-forming at all locations, although it was just shy of reaching the 0.5 threshold for group-forming at PNG (Fig 3). Other species also varied due to nonbreeding subordinates being accepted into a coral depending on location. However, some species that were originally defined as group-forming switched to pair-forming post-disturbances (as subordinates co-habited less often post-disturbances), suggesting that group-forming species have moderate degrees of sociality (Fig 3). *Gobiodon citrinus* was the only species to remain group-forming regardless of location or disturbance and to have the most subordinates in groups (highest sociality indices). However, this species was rarely encountered and only found in sufficient numbers for sociality index calculation post-disturbance at OTI. *Gobiodon fuscoruber* was initially group-forming at all locations albeit with a lower sociality index than other group-forming species, except at PNG where it was defined as pair-forming. At OTI, this species remained group-forming post-disturbance with little change to their sociality index. At LI, this species was too rare post-disturbances for analysis to be conducted for those years. *Gobiodon rivulatus* was another species that had the highest sociality index with many subordinates co-habiting at LI pre-disturbance, but it became exclusively pair-forming without subordinates at LI post-disturbances. At PNG, this species was defined as pair forming, just falling shy of a 0.5 sociality index. However, the species was exclusively pair-forming at OTI pre-disturbances, and instead occasionally accepted subordinates at OTI post-disturbances (Fig 3).

**Fig 3.**
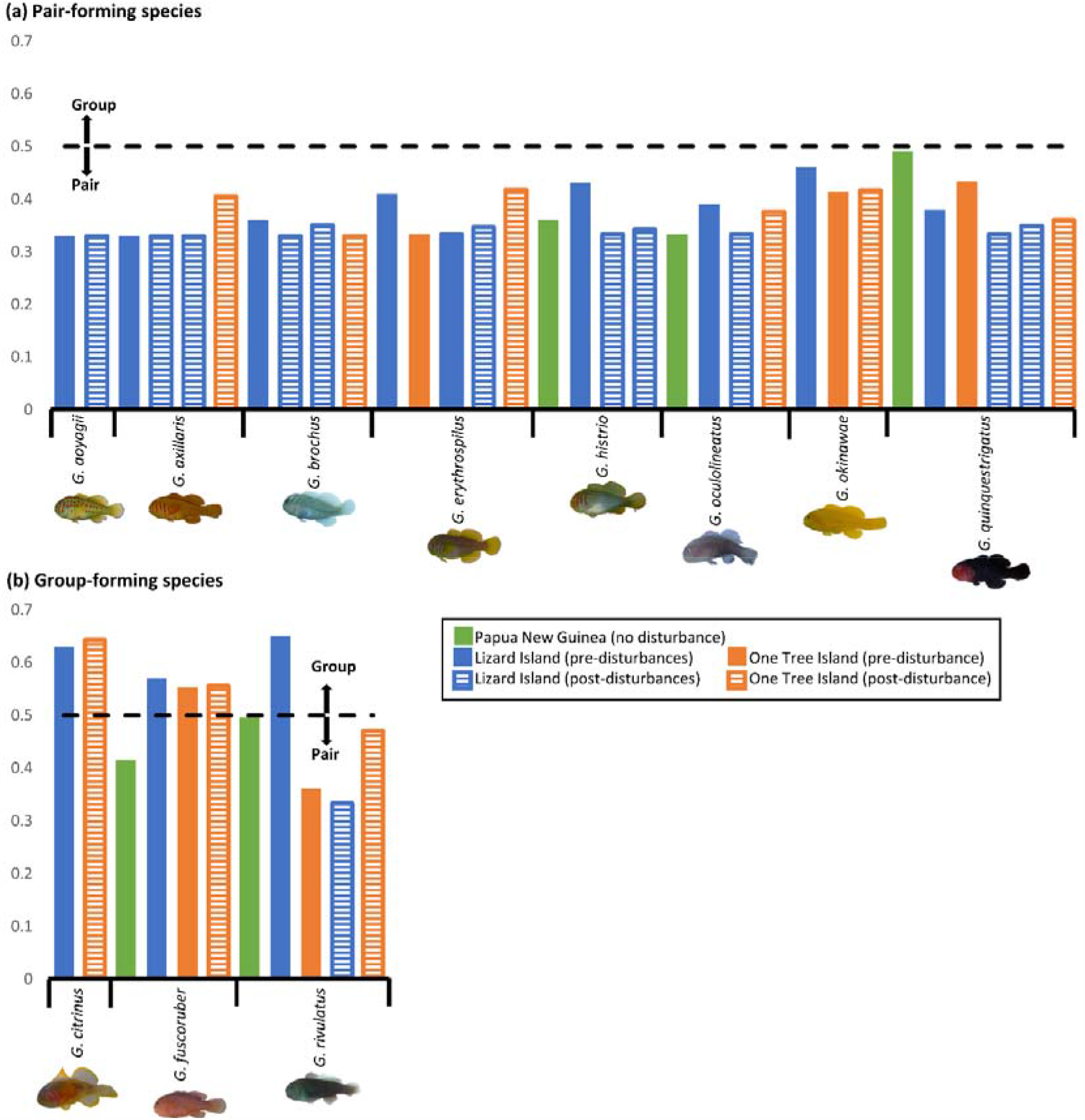
Sociality index of each species at different locations including repeat visits pre- and post-disturbance(s).

### 4.3. Social Plasticity: Group Size – Size of the Dominant – Coral Size

We had sufficient sample size to compare 2 group-forming species (i.e. *G. fuscoruber* and *G. rivulatus*) at 2 locations (LI, OTI). We investigated the relationship between group size, size of the dominant individual, and coral size (Fig 4). For both species, group size was positively related to coral size (Suppl. Tab 2, p < 0.01), but was not related to the size of the dominant individual or location (p > 0.40). For *G. rivulatus*, the size of the dominant individual was positively related to coral size for both species (p < 0.05), and to group size and location (p = 0.03, p < 0.01, respectively). For *G. fuscoruber*, the size of the dominant was not related to group size or location (p > 0.36). for both species, Coral size was positively related to group size and the size of the dominant (p < 0.01) but was not related to location (p > 0.14). There was no interaction between any of the variables for each analysis (p > 0.27). Note: no analyses were completed to compare these size relationships pre-versus post-disturbance as colonies were primarily made up of pairs at LI post-disturbance and no colonies were collected at OTI post-disturbance.

**Fig 4.**
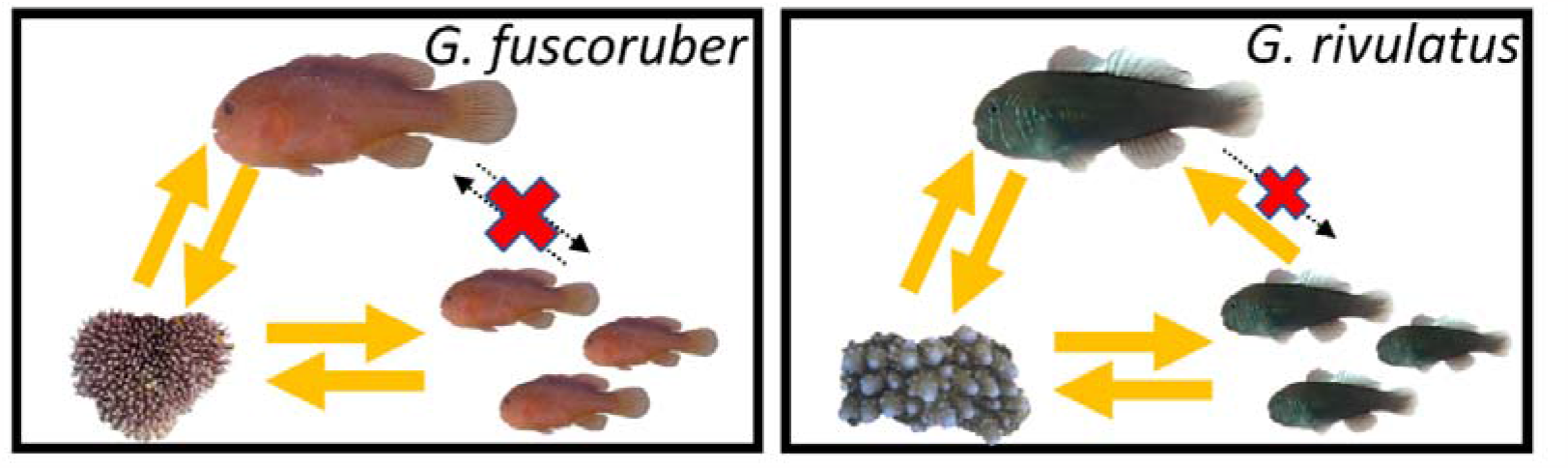
Social plasticity in group size, size of dominant, and coral size for group-forming *Gobiodon* gobies and their *Acropora* coral hosts. Yellow arrows identify significant effect (p < 0.05), and crossed out dashed lines represent no significant effect (p ≥ 0.05).

### 4.4. Within-group plasticity: Size Ratios

We compared the size ratios between rank 1 and rank 2 (i.e. rankstep 1) for six species (*G. erythrospilus, G. fuscoruber, G. histrio, G. oculolineatus, G. quinquestrigatus*, and *G. rivulatus*) that were found at multiple locations with sufficient sample size. Mean size ratio for rankstep 1 ranged between 0.88 and 0.94 ± 0.01-0.02 among all species (Fig 5). Size ratios for rankstep 1 were not related to coral size (Suppl. Tab 3, p = 0.94), group size (p = 0.09), species (p = 0.15) or location (p = 0.52), and there was no interaction between any predictors (p = 0.24). Since there was no effect of location, we then included a seventh species, *G. brochus*, that was only found at one location (LI). Including *G. brochus* did not change the outcome of the model with size ratios for rankstep 1 being unrelated to coral size (p = 0.21), group size (p = 0.25), and species (p = 0.12).

**Fig 5.**
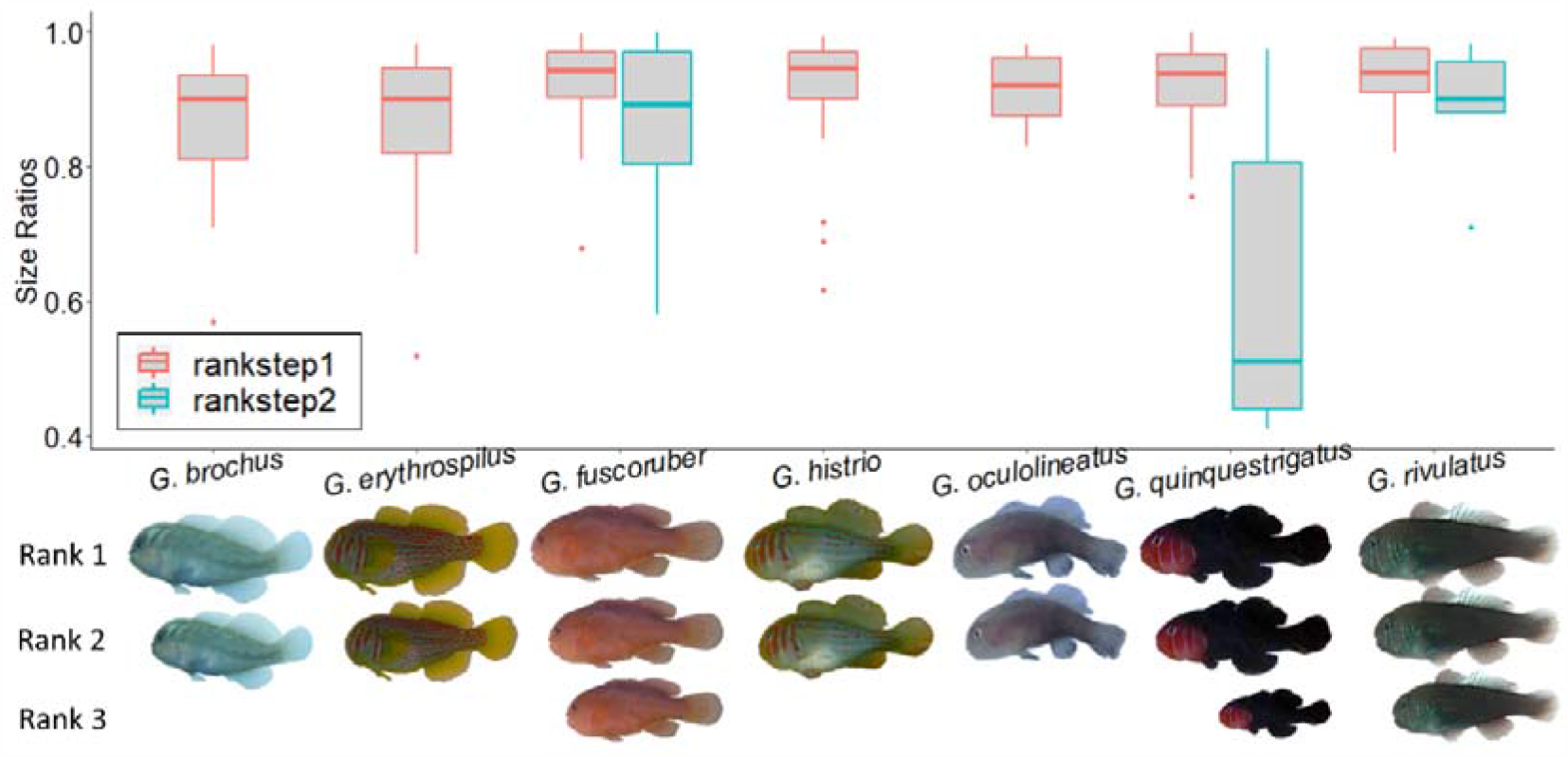
Distribution of size ratios between ranks 1 & 2 (rankstep1), and ranks 2 & 3 (rankstep2) of single species colonies of *Gobiodon* species. Note: there is no rankstep2 data for *G. brochus, G. erythrospilus, G. histrio*, and *G. oculolineatus* due to insufficient data; and the size differences between ranks for each species are shown with pictures that are illustrated to scale based on rankstep means.

For size ratios between rank 2 (second breeder) and rank 3 (first nonbreeder) (i.e. rankstep 2), there were insufficient colonies with rank 3 individuals for four of the seven species (*G. brochus, G. erythrospilus, G. histrio*, and *G. oculolineatus*), so these species were excluded. Further, we pooled the size ratios for rankstep 2 for the other 3 species among locations, because there were not enough samples per location and location did not affect size ratios for rankstep 1. The size ratio for rankstep 2 was slightly lower than rankstep 1 for most species (Fig 5). Size ratios for rankstep 2 were related to coral size (p = 0.003), group size (p = 0.003), and species (p = 0.05). Specifically, there was a positive relationship between size ratios, coral size and group size. Rank 3 tended to be much smaller for *G. quinquestrigatus* (rankstep 2 mean = 0.63 ± 0.11) than other species (rankstep 2 mean ranging from 0.85 to 0.90 ± 0.03-0.08). for *G. quinquestrigatus*, the smaller rank 3 individuals suggests that the species is primarily pair-forming, but that breeders will tolerate nonbreeders occasionally if they are far smaller in size (Fig 5).

We revisited LI and PNG in consecutive years (LI2020 and LI2021, PNG2018 and PNG2019), and calculated the size ratio for rankstep 1 if both dominant individuals tagged in the first trip were still present in the following trip. The size ratios for rankstep 1 were related to coral size (p = 0.02), but not to group size (p = 0.76), species (0.30), location (p = 0.37), nor year (p = 0.09), and there were no interactions (p > 0.07). The time between visits at LI was one year compared to only six months at PNG, and yet there was no effect of location or interaction with year on the size ratios. Although the effect of year was not significant, there is a trend for rank 1 and rank 2 individuals to converge in size overtime (Suppl Fig 2).

When comparing the size ratio of rankstep1 pre- and post-disturbances at LI, we only had sufficient sample sizes for *G. brochus, G. erythrospilus, G. histrio*, and *G. quinquestrigatus*. The size ratio of rankstep 1 was related to coral size (p < 0.01), but not to group size (p = 0.06), species (p = 0.19), or pre-vs. post-disturbance (p = 0.29), and there was no interaction (p = 0.20).

### 4.5. Within-group Plasticity: Sex Dominance Between Breeding Partners

Sex dominance was only identified during trips to LI in 2020 and 2021. We compared sex dominance in goby colonies only if both dominant individuals tagged in 2020 were still present in 2021. There were five goby species found in high enough abundance to determine whether sex dominance existed for rank 1. In 2020, 120 colonies were identified for sex dominance, and 42 colonies were revisited in 2021. From both years combined, the sex ratio between rank 1 females and rank 1 males was 1:0.7 which differed significantly from unity (Suppl. Tab 3, p = 0.02). There was also a difference among years (p < 0.01): in 2020, the ratio of female to male rank 1 was 1:1.05 among species (Suppl Fig 3), whereas in the same colonies in 2021, females often outgrew males and the sex ratio was 1:0.36 female to male rank 1 individuals for all species (Suppl Fig 3). The male never outgrew the female in any colonies (Suppl Fig 3). There was no difference in the ratio of female to male rank 1 individuals among species (p = 0.30) and no interaction between species and year (p = 0.29).

### 4.6. Mixed Species Colonies: Social Structure and Composition

Although we did not sex individuals to confirm they were reproductively active, we did find two nests containing eggs within the same coral on more than one occasion. Each nest was being guarded by a specific same-species pair, suggesting that sharing of nest guarding was not occurring and there was likely no hybridization. It is also important to note that no mixed species colonies were collected post-disturbance at any of the locations, therefore no pre-versus post-disturbance analyses were completed for mixed species analyses.

When quantifying the size-based hierarchy within mixed groups, we found that different species were intermixed within the ranks just under 50% of the time with no difference among locations (intermixed e.g. rank 1,3,5 were species A and rank 2,4,6,7 were species B, Suppl. Tab 4, p = 0.91, Fig 6A, Suppl Fig 4). The rank 1 individual within mixed groups was generally the larger-bodied species (as defined by Hing et al. 2019) approximately 75% of the time with no pattern among locations (p = 0.93, Fig 6B). Although different species were intermixed within ranks, we propose that individuals still queue for a breeding position within their own species as eggs were guarded by pairs of the same species.

**Fig 6.**
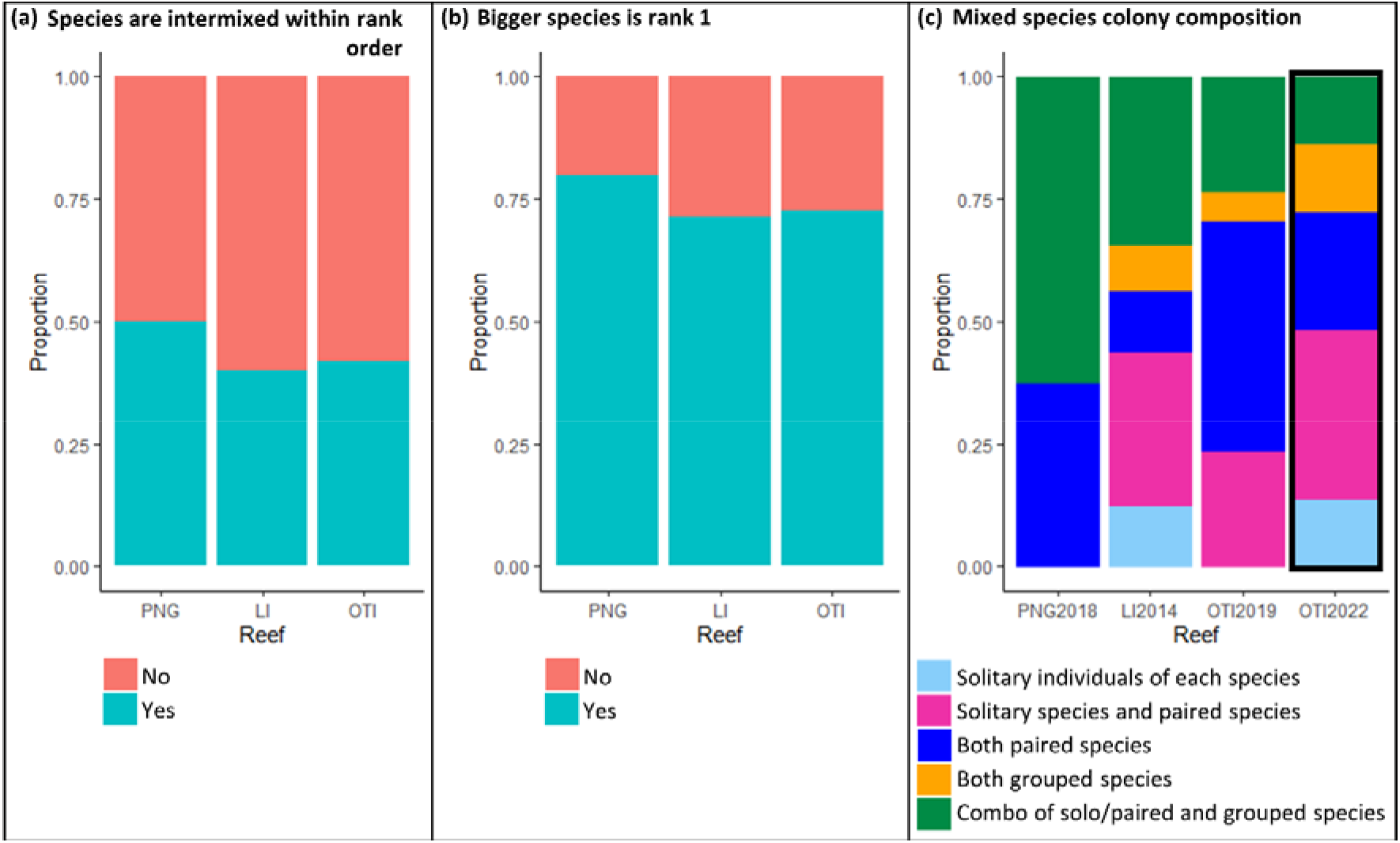
Proportion of intermixed ranks **a** and larger-bodied species as rank 1 **b** within size-based hierarchies of mixed species colonies of *Gobiodon* and their grouping composition **c**. PNG = Papua New Guinea; LI = Lizard Island; OTI = One Tree Island; year after the location label is the year sampled; data outlined in thick black line was taken post-disturbance while all other data was taken pre-disturbance.

When we calculated the size ratios between each rank within mixed species colonies, there were sufficient large groups to compare ranksteps 1-8 (i.e. from rank 1 down to rank 9). The size ratio of each rankstep in mixed species colonies differed by coral size (Suppl. Tab 2, p < 0.01), group size (p = 0.02), but not by rankstep (p = 0.10) or location (p = 0.11). There was no interaction between any of the variables. There was a positive relationship between size ratios and coral sizes as well as group sizes. This means that ranks were more similar in size in larger corals and in bigger groups. We found that when size ratios were separated per species, size ratios within mixed species colonies were smaller on average (0.88 ± 0.01) than those for that same species in single species colonies (0.91 ± 0.01, p < 0.01). The smaller size ratios in mixed species colonies means that group sizes in mixed colonies may be as large as in single species groups due to rules of the size hierarchy with respect to group size (Buston & Cant 2006; Wong 2011). We then compared size ratios of mixed species colonies, regardless of species, to single species colonies and found no difference between mixed or single species colonies (p = 0.22, Fig 5& Suppl Fig 4).

Pre-disturbances, mixed species colonies were composed of solitary, pair-forming and/or group-forming species with no difference in proportion among locations (Suppl. Tab 4, p = 0.69, Fig 6C). There was also no difference in mixed species composition pre- or post-disturbance at OTI (p = 0.58, Fig 6C). Note, not enough mixed species colonies were found at LI post-disturbance, therefore LI was not compared for disturbance effect. Mixed species colonies were primarily made up of two species (89%), followed by three species (10%), and there was only a single colony of four species (1%, Suppl. Tab 5). Every *Gobiodon* species observed was found in a mixed species colony at least at one time point (Suppl. Tab 5). However, the most common mixed species colonies were made up of *G. fuscoruber-G. quinquestrigatus* colonies (23%), followed by *G. fuscoruber-G. rivulatus* colonies (10%), and then *G. oculolineatus-G. quinquestrigatus* colonies (9%, Suppl. Tab 5). The single most common species in mixed species colonies was *G. fuscoruber* (55%), followed by *G. rivulatus* (43%), and *G. quinquestrigatus* (41%, Suppl. Tab 4). The following species were found with similar proportions within mixed species and single species colonies: *G. citrinus, G. fuscoruber, G. oculolineatus*, and *G. diabolensis* (Hildebrandt *et al*., in prep); all other species observed occurred more often in single species colonies than in mixed species colonies (Suppl. Tab 5).

## 5. Discussion

Following our sociality framework, we investigated to what extent ecological and social factors affected the four categories of social variation in coral-dwelling gobies in the *Gobiodon* genus. We chose both large-scale ecological factors, namely location and disturbance regime, and small-scale ecological factors, namely habitat characteristics like habitat size, as well as social factors, namely body size of the largest group member. Each category of variation outlined in the framework (i.e. forms of sociality, degree of sociality, social plasticity, and within-group plasticity) guided our assessment of the relevant ecological and social factors. We found that location and disturbance regimes played a substantial role in the forms of sociality exhibited within the genus and the degree of sociality exhibited by individual species, with species tending away from group-forming under high disturbance regimes. In contrast, social plasticity and within-group plasticity were not directly affected by these large-scale factors but were indirectly affected by small-scale factors like changes to coral size, which decreased following disturbances. Based on these findings, we infer that societies of coral-dwelling gobies have an extremely poor outlook in terms of persistence when facing climatic disturbances. Accordingly, this framework allowed us to identify the impacts of multiple ecological factors on animal societies over different scales.

With respect to the form of sociality, studying multiple goby species within the *Gobiodon* genus enabled us to investigate how the form of sociality within the whole genus was affected by small and large scale ecological factors. Coral sizes affected the form of sociality, with a shift from solitary to pairs and groups as coral sizes increased. In addition, location was also a key predictor. In the northern location at Kimbe Bay, Papua New Guinea, gobies tended to form pairs; in the central reef location at Lizard Island, Australia gobies tended to form single species groups; and in the southern location at One Tree Island, Australia, gobies tended to form mixed species colonies. This gradient may indicate a latitudinal shift in social systems, as seen in ground-nesting bees (Dew *et al*. 2018) and birds (Arnold & Owens 1998). Reef type may potentially explain such differences between locations; for example, the movement of goby larvae may be limited in a lagoonal reef like at One Tree Island and prompt the formation of mixed species colonies in order for more individuals to populate an area while reducing the potential for inbreeding (Selwyn *et al*. 2016). It should be noted though that we did not sample at multiple locations at each latitude, hence limiting our ability to draw conclusions as regarding the major underlying causes of this latitudinal variation.

Additionally, disturbance regime was a strong predictor of forms of sociality, with high disturbance regimes reducing the propensity for group-living as gobies were found either living solitary or in pairs after these disturbances. After moderate disturbance regimes, gobies were also primarily living solitary and less often in pairs, but the same proportion of groups was still found compared to pre-disturbances. Finding many solitary gobies is a cause for concern as pairs are needed for breeding. Such a loss in sociality due to disturbance is likely due to the extreme decline in populations of gobies following particularly extreme events (Froehlich *et al*. 2021; Hing *et al*. 2018). The increased occurrence of solitary living could be attributed to a reduction in a) space and shelter due to corals becoming damaged and/or b) food resources which the corals provide. A similar result was reported for passerine birds when habitat size was reduced after disturbance (Lantz & Karubian 2017) and in butterflyfishes when food resources were reduced after disturbance (Thompson *et al*. 2019). Ecological factors such as environmental disturbances are therefore important predictors for the form of sociality within the genus of *Gobiodon*.

While location and disturbance regime were observed to reduce the degree of sociality for group-forming species, they did not change the degree of sociality for pair-forming species. These latter species exhibited low degrees of sociality and tended to live in pairs (0.33-0.49) regardless of location or disturbance regime. This study further provides support that these species are generally pair-forming as was also determined by Hing *et al*. (2018, 2019). Interestingly, although these pair-forming species primarily live in pairs, some accepted nonbreeding subordinates during periods of low disturbances, but did not accept any nonbreeders during high disturbance regimes. Furthermore, some group-forming species (Hing *et al*. 2018, 2019) displayed moderate degrees of sociality (0.33-0.65) that fluctuated between group- or pair-forming depending on location and disturbance regime, and these patterns were not always similar among species. Typically, the degrees of sociality fluctuated post-disturbance depending on the species surveyed. The two group-living species, *G. citrinus* and *G. rivulatus* had the highest degrees of sociality at one location (i.e. LI, with most subordinates in a group), and either continued occurring in groups with many subordinates (*G. citrinus*), or became pair-forming post-disturbances (*G. rivulatus*). The third group-living species, *G. fuscoruber*, remained as group-living at most locations (LI, OTI) pre-disturbances and after low disturbances, although with fewer subordinates post-disturbances. However, it is important to note that *G. citrinus* and *G. fuscoruber* disappeared after extreme disturbances at Lizard Island (Froehlich *et al*. 2021). Different species appear to have different responses to disturbances in terms of their sociality index, and we need further work to understand the fitness consequences of these species-specific differences.

The life insurer hypothesis (Queller & Strassmann 1998) states that cooperative and social groups enjoy a competitive advantage in challenging habitats, hence why sociality has evolved. In support of this, several studies have demonstrated that species have a higher chance of survival in challenging environments by living socially instead of paired or solitary due to benefits of resource acquisition, brood care, and predator protection (Duffy & Macdonald 2010; Firman *et al*. 2020; Queller & Strassmann 1998; Rubenstein & Lovette 2007) (Queller & Strassmann 1998),. Our results did not provide support for the life insurer hypothesis of sociality, the goby species had lower degrees of sociality in challenging environments (i.e. high disturbances). In comparison, In comparison, based on global study, many birds evolved social living as a strategy to ensure survival in environments that are constantly fluctuating and challenging (Jetz & Rubenstein 2011). Similarly, naked molerats have some of the highest degrees of sociality and are strictly eusocial, like *Heterocephalus glaber* (0.95) and *Fukomys damarensis* (0.80-91) (Avilés & Harwood 2012), and they live socially due to challenging environments that fluctuate substantially in rainfall (Faulkes *et al*. 1997). By comparison, gobies exhibit low to moderate degrees of sociality, and ecological conditions play a large role in their grouping tendencies. Our study suggests that gobies likely evolved social living behaviour in stable environments, as seen in hornbills (Gonzalez *et al*. 2013). In stable environments, corals can grow larger, and more gobies can reside within a coral and reap the benefits of sociality. When conditions deteriorate and corals become smaller, group-living is no longer possible hence why gobies switch to living with fewer subordinates and primarily in pairs in challenging environments.

When addressing variations in sociality at smaller scales, we found that the group sizes of group-forming gobies were plastic with respect to habitat size, but not i location. This demonstrates that coral size is a key limiting resource influencing sociality, as gobies were in smaller groups when corals became smaller after climatic disturbances (Froehlich *et al*. 2021; Hing *et al*. 2018, 2019; Madin *et al*. 2018). For the most social goby species studied at all locations, *G. fuscoruber*, the coral size influenced the size of the dominant individual and group size, but the size of the dominant was not influenced by the group size. This suggests that social constraints on group size, namely the size of the largest dominant individual, have less of an influence on group size than ecological factors like coral size. On the other hand, group sizes of *G. rivulatus* affected the size of the dominant individual, but not vice versa. Location had little impact on any of these relationships for either species. In contrast, all three variables (group size, habitat size, and size of dominant) were positively related to each other for other social fishes like *P. xanthosoma* and *A. percula*, suggesting strong social plasticity based on habitat size and social context (Buston & Cant 2006; Wong, 2011; Rueger et al. 2021). Social plasticity therefore appears to vary depending on the species and its ecology, and such variation highlights that integrating large-scale factors into investigations alongside small-scale factors can provide important insights into social plasticity.

In terms of within-group plasticity, we found that size ratios and sex dominance ratios of gobies were not directly affected by large scale ecological factors like location and disturbance. We found that size hierarchies of *Gobiodon* are similar to those of *Paragobiodon* (Wong et al. 2007, 2008); in a goby colony the two dominant individuals are slightly different in size (1:0.88 to 1:0.94) regardless of species or location. Breeding partners likely converge in size over time to maximize reproductive output (Munday *et al*. 2006), although not for all cases. Coral size influenced the size ratios between breeding partners for some but not all analyses, suggesting that size convergence may not be beneficial in all circumstances. We found that although males are often the bigger individual at first, females will outgrow males more than half of the time, owing to their growth rate advantage (Munday *et al*. 2006; Nakashima *et al*. 1996). Initially a bigger male allows for better paternal care and offspring success in the first breeding year, but then a bigger female allows for more offspring in a single egg clutch (Nakashima *et al*. 1996). Gobies also have bi-directional sex change which allows either individual to change sex if their mate dies and they find a new partner (Munday *et al*. 1998; Nakashima *et al*. 1996; Sunobe *et al*. 2017). This suggests that while a bigger female is advantageous in the long run, groups are not strictly matriarchal like those in the anemonefish *A. percula* (Buston & Wong 2014; Rueger *et al*. 2021a; Wong & Buston 2013).

When considering variation in size ratios in colonies, specifically between rank 2 and 3, we found that their size ratio is slightly smaller (0.85 to 0.9) than that between the breeding individuals (rank 1 and rank 2) for most species i.e. there is a larger size gap between rank 2 and 3 than between rank 1 and 2. However, rank 3 nonbreeders for *G. quinquestrigatus* considerably smaller than the rank 2 individuals (0.64). This is not entirely surprising as *G. quinquestrigatus* was living primarily in pairs, suggesting limited tolerance of breeders for any nonbreeder. Regardless of species, the two breeders (rank 1 and 2) were closer in size than the first nonbreeder (rank3) was to the closest breeder (rank 2). This is expected as breeders converge in size for reproductive benefits (Kuwamura *et al*. 1993; Munday *et al*. 2006), whereas nonbreeders regulate their sizes to be tolerated\ by breeders and avoid eviction (Wong *et al*. 2007). This provides evidence that *Gobiodon* gobies cooperate within size-based hierarchies, as seen in *P. xanthosomus* (Kuwamura *et al*. 1993; Wong *et al*. 2007). Size ratios between rank 2 and 3 were affected by coral size with rank 3 being more similar in size as the rank 2 breeder in larger corals for pair-forming species. This suggests that larger corals provide nonbreeders with more opportunities to grow larger and be more tolerated by breeders. Therefore, living in groups may be costly for nonbreeders from pair-forming species as they must remain far smaller than breeders despite coral size, making group-living only potentially advantageous in large corals (Hing *et al*. 2019; Rueger *et al*. 2021a). However, for strictly group-forming species, there was little effect of coral size, suggesting that breeders are more tolerant of nonbreeders and appear to allow nonbreeders to grow larger regardless of coral size (Rueger *et al*. 2021a).

When investigating the within-group composition of mixed colonies, we found that that different species were often interspersed in ranks within the hierarchy. Interestingly though, the size ratios between ranks remained the same regardless of which species were adjacent in ranks, and regardless of location. Instead, individuals can grow larger in larger corals as there is likely more space to avoid aggression from higher ranks. With a larger coral, dominant individuals can grow larger, thus allowing additional subordinate individuals to fit within the size-based hierarchy. Individuals were also closer in size in larger corals, suggesting that larger corals may reduce conflict among individuals. When factoring in the clear size differences between goby species, with some species growing larger on average than others (Hing *et al*. 2019), we found that the larger-bodied species tended to occupy the rank 1 position (i.e. largest individual) in mixed species colonies, regardless of location. Accordingly, this suggests that *Gobiodon* cooperate within size-based hierarchies in both single species colonies and mixed species colonies alike.

There was no clear trend for whether mixed species colonies were composed of only pair-forming individuals, groups, or a combination of both, regardless of location or disturbance regime. However, some species were more often found in mixed species colonies than others. For example, *G. fuscoruber, G. quinquestrigatus* and *G. rivulatus* were more often found in mixed species colonies than other species, suggesting they obtain some advantage to living in mixed colonies (Ellis & Good 2006). By far the most common mixed species colony was composed of *G. fuscoruber* and *G. quinquestrigatus* One potential advantage of living with congeners may be that individuals can reach breeding status quicker (as there may be fewer conspecifics queuing for breeding status) while still receiving synergistic benefits of living in a larger group (Rueger *et al*. 2021a). Indeed, we found evidence for separate breeding queues for each species within a colony. On multiple occasions mixed species colonies had two egg clutches within the coral - one guarded by a pair from one species and another guarded by a pair from another species (pers obs). Gobies in mixed species colonies can reap the various benefits of living in big groups (e.g. improved territory defence, improved coral growth, improved survival and growth rates) whilst not necessarily decreasing their likelihood of territory inheritance and securing reproduction (Goodale *et al*. 2017; Rueger *et al*. 2021a). In order to maintain cooperation, gobies in mixed species colonies regulate growth in size-based hierarchies just like in single species colonies. Future studies comparing egg clutch sizes, rates of territory defense, long term growth rates, and survivorship among single species and mixed species colonies would be important in identifying the benefits of living in mixed species colonies.

In each of the four categories of variation, we found direct and indirect impacts of climatic disturbances, suggesting an extremely high loss of sociality (Fig 1). The form of sociality and degrees of sociality were each drastically lower after high disturbance regimes. Social plasticity and within-group plasticity were not directly affected by disturbances, but instead were indirectly affected via a decrease in coral size. Since disturbances drastically diminish the sizes of available corals (Froehlich *et al*. 2021; Hing *et al*. 2018, 2019; Madin *et al*. 2018), social plasticity and within-group sociality are indirectly lost to disturbances. Accordingly, each category of variation in coral-dwelling goby societies is facing high loss to disturbances. Given that living in groups can increase individual fitness and survival (Booth 1995; East & Hofer 2010; Gil *et al*. 2017; Jordan *et al*. 2009; Komdeur & Ma 2021; Strauss & Holekamp 2019), these findings suggest that the large-scale population losses observed in coral-dwelling gobies after environmental disturbances (Froehlich *et al*. 2021, 2023) is at least in part due to a loss of sociality at multiple levels.

By quantifying the four categories of variation, the sociality framework introduced here provides a flexible yet robust assessment of social organisation for animal societies along different scales of ecological and social factors. Depending on the factors of interest, each category of variation can be quantified at a defined spatial and temporal scale. The framework can identify limiting resources that will play important roles in the formation and maintenance of animal societies. The framework is particularly useful as it requires only monitoring of group sizes, measures of cooperation, e.g. size and sex of individuals within groups, and measures of ecological and social factors of interest, e.g. habitat size and proximity to other groups, without requiring manipulative experimentation. The categories of variation (i.e. forms of sociality, degree of sociality, social plasticity, and within-group plasticity) as well as the social and ecological factors can be easily adapted to the life history, cooperation, and ecology of the social taxon (e.g. Fig 1). The framework can be adapted for any species and many different factors, including larger-scale ones like spatiotemporal and disturbance factors, thus making observational data a powerful tool for modeling the social organisation and plasticity of many taxa into the future. By assessing how each category of variation is affected by ecological factors, the metrics can then be integrated to identify the outlook of social maintenance in the taxon studied.

## Data Availability Statement

Data and R scripts will be available at knb.ecoinformatics.org.

## Acknowledgements

The field trips were conducted with respect for the traditional owners past and present of the Kilu and Tamare communities of Kimbe Bay in Papua New Guinea, Dingaal Aboriginal Traditional Owners of Lizard Island, and Gurang, Gooreng, Gooreng, Bailai and Bunda Traditional Owners of One Tree Island. We greatly appreciate the time and effort that our field helpers have put towards this study: Kylie Brown, Karen Hing, Grant Cameron, Anna Scott, Rituraj Sharma, Theresa Rueger, Ana Gaisinier, Sabrina Velasco, Nelson and Jerry Sikatura. We are grateful to the station management and facilities at the Mahonia Na Dari Research and Conservation Centre of Kimbe Bay Papua New Guinea, Lizard Island Research Station, and One Tree Island Research Station, and particularly Anne Hoggett, Lyle Vail, Peter and Jane Miller, Somei Jonda, Ainsley and Paul Carin, Heinrich Breuer and Ruby Holmes for their assistance with our field trips. We would like to thank Jackie Wolstenholme and Zoe Richards for help with coral ID confirmation.

## Ethics approval

The study followed relevant guidelines and regulations, including ARRIVE guidelines, and was conducted under the animal ethics protocols of the University of Wollongong (protocols: AE1404, AE1725, and AE2117). Permit approvals were acquired from the Great Barrier Reef Marine Park Authority (permits: G13/36197.1, G15/37533.1, G18/41020.1) and from the Papua New Guinea Research Visa Permit Office (permit: AA654347).

## Author contributions

CF: planned study, sought funding, collected data, analysed data, wrote and edited the manuscript; SH: collected data, analysed data, edited the manuscript; OK: planned study, collected data, edited the manuscript; MH: collected data, edited manuscript; CH: collected data, edited manuscript; JS: collected data; MW: planned study, sought funding, collected data, edited the manuscript.

