## Supplementary information for "Multi-level framework to assess social variation in response to ecological and social factors: modeled with coral gobies"

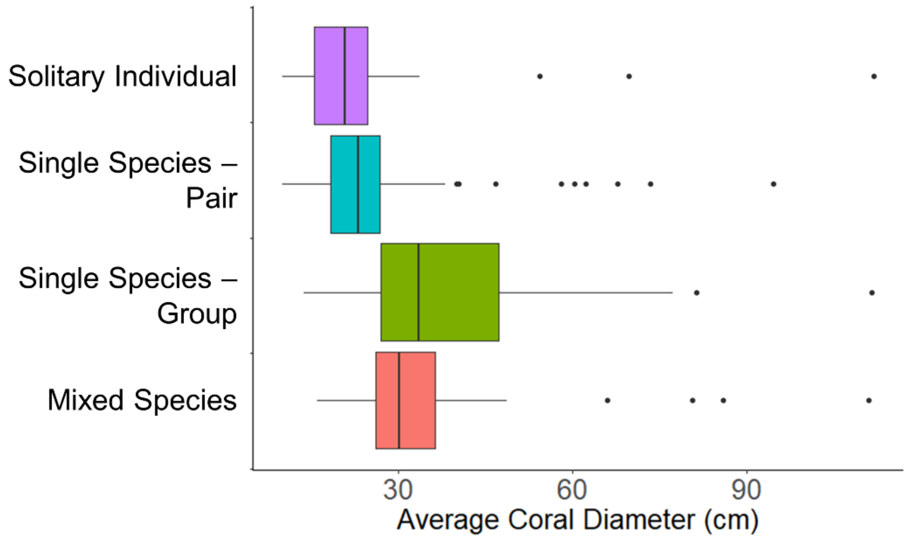


Suppl Fig 1. Relationship between form of sociality and coral size.

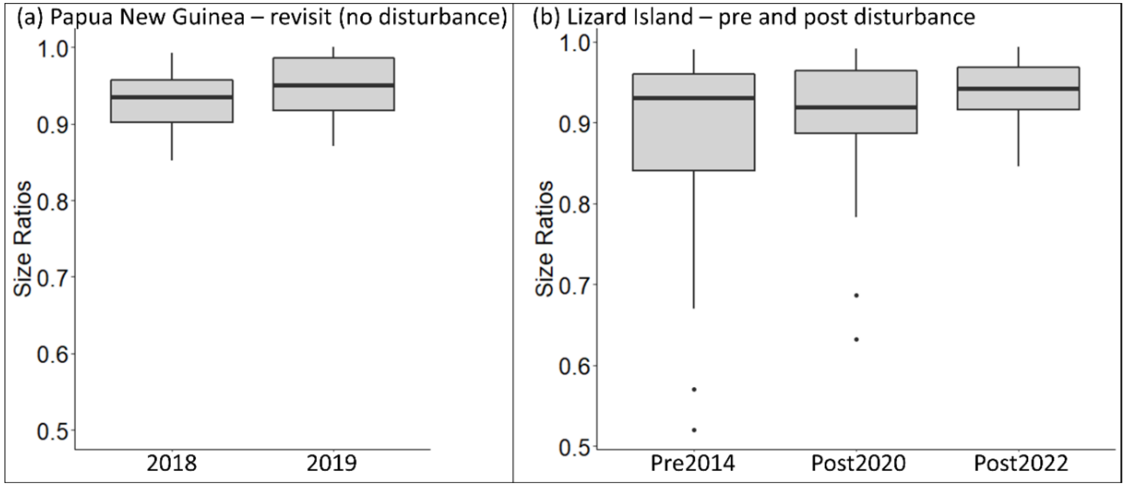


Suppl Fig 2. Size ratio between rank 1 and rank 2 individuals of *Gobiodon* species within single species colonies that were revisited at **a** Papua New Guinea (Sep-Nov 2018 and May-June 2019) and **b** Lizard Island before and after disturbances and follow up visit (Jan-Feb 2014, Jan-Mar 2020 and Jan-Mar-2021)

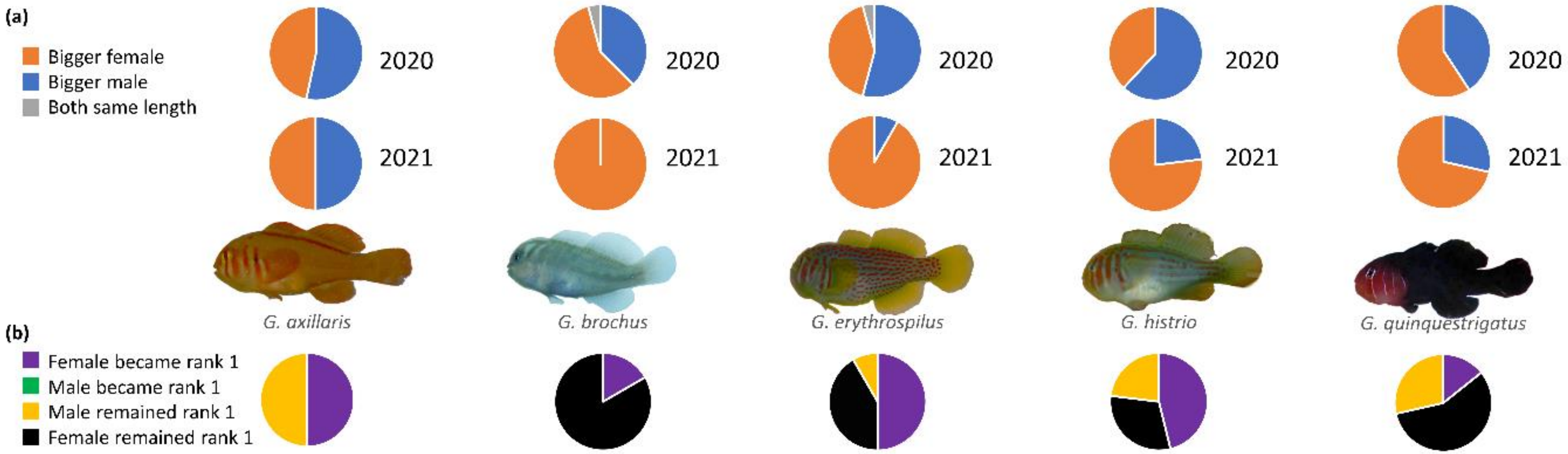


Suppl Fig 3. **a** Sex dominance of species visited at Lizard Island in 2020 vs. 2021, and **b** revisited the following year to see whether any sex outgrew the other in 2021. Note: no male outgrew the female in any goby colony (green).

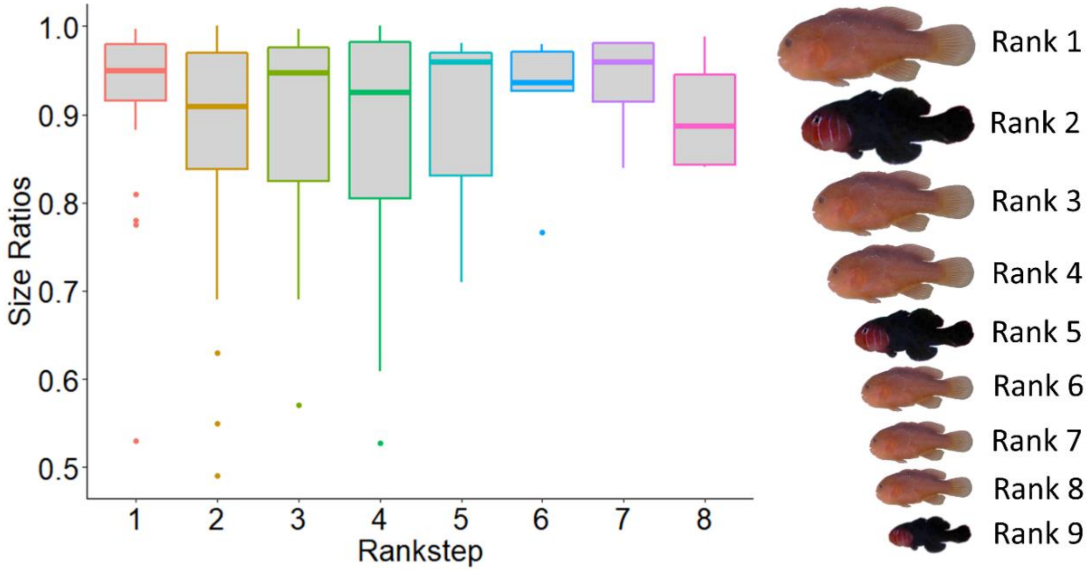


Suppl Fig 4. Size ratios between each rankstep within the size-based hierarchies of mixed species colonies of *Gobiodon* gobies. Note: rankstep i = ratio between $\text{rank}_{(i)}$ and $\text{rank}_{(i+1)}$ individuals; the size differences between ranks are shown with pictures that are illustrated to scale based on rankstep means.

Supplementary Table 1. Statistical output of form of sociality analyses. VGLM = multinomial logistic regression model

| Response Variable | Model | Predictor variable | Factor Type | df | Model Statistic | Test-value | p-value | R-squared statistic | Test value |
|---|---|---|---|---|---|---|---|---|---|
| Form of Sociality pre-disturbances only (categorical variable: 4 levels) | VGLM | average coral diameter | covariable | 3 | χ2 (chi-squared) | 55.73 | < 0.0001* | McFadden | 0.102 |
| | | location | fixed | 6 | | 21.88 | 0.0013* | Cox and Snell (ML) | 0.206 |
| | | | | | Sample size | n = 529 | | Nagelkerke | 0.23 |
| Form of Sociality pre and post-disturbances (categorical variable: 4 levels) | VGLM | average coral diameter | covariable | 3 | χ2 (chi-squared) | 46.95 | < 0.0001* | McFadden | 0.131 |
| | | location | fixed | 3 | | 24.05 | < 0.0001* | Cox and Snell (ML) | 0.245 |
| | | pre/post | fixed | 3 | | 84.00 | < 0.0001* | Nagelkerke | 0.277 |
| | | location x pre/post | fixed | 3 | | 42.00 | < 0.0001* | | |
| | | | | | Sample size | n = 1239 | | | |
| Form of Sociality x year post-disturbance at Lizard Island (categorical variable: 4 levels) | VGLM | average coral diameter | covariable | 3 | χ2 (chi-squared) | 65.25 | < 0.0001* | McFadden | 0.086 |
| | | year | fixed | 3 | | 21.29 | < 0.0001* | Cox and Snell (ML) | 0.146 |
| | | | | | | | | Nagelkerke | 0.174 |
| | | | | | Sample size | n = 791 | | | |

Supplementary Table 2. Statistical output of social plasticity analysis. GLM = generalized linear model, LM = linear model.

| Response Variable | Model | Predictor variable | Factor Type | df | Model Statistic | Test-value | p-value | R-squared statistic | Test value |
|---|---|---|---|---|---|---|---|---|---|
| Group Size | GLM | average coral diameter | covariable | 1 | χ2 (chi-squared) | 8.78 | 0.006* | marginal | 0.267 |
| for *G. rivulatus* | | size of dominant individual | covariable | 1 | | 7.66 | 0.29 | | |
| (includes all coral species) | | reef | factor | 2 | | 5.82 | 0.40 | | |
| | | coral diam x dominant size | interaction | 1 | | 5.79 | 0.85 | | |
| | | | | | Sample size | n = 37 | | | |
| | | | | | Outliers removed | n = 1 | | | |
| Size of Dominant Individual | LM | average coral diameter | covariable | 1 | F-value | 4.37 | 0.045* | marginal | 0.423 |
| for *G. rivulatus* | | group size | covariable | 1 | | 5.10 | 0.03* | | |
| log-transformed | | location | factor | 2 | | 5.93 | 0.007* | | |
| (includes all coral species) | | coral diam x group size | interaction | 1 | | 0.62 | 0.44 | | |
| | | | | | Sample size | n = 37 | | | |
| | | | | | Outliers removed | n = 1 | | | |
| Coral Diameter | LM | size of dominant individual | covariable | 1 | F-value | 7.84 | 0.009* | marginal | 0.522 |
| for *G. rivulatus* | | group size | covariable | 1 | | 21.76 | <0.0001* | | |
| log-transformed | | location | factor | 2 | | 2.13 | 0.14 | | |
| (includes all coral species) | | dominant size x group size | interaction | 1 | | 0.0008 | 0.98 | | |
| | | | | | Sample size | n = 37 | | | |
| | | | | | Outliers removed | n = 0 | | | |
| Group Size | GLM | average coral diameter | covariable | 1 | χ2 (chi-squared) | 3.91 | 0.0002* | marginal | 0.412 |
| for *G. fuscoruber* | | size of dominant individual | covariable | 1 | | 3.68 | 0.63 | | |
| (includes all coral species) | | location | factor | 2 | | 3.60 | 0.96 | | |
| | | coral diam x dominant size | interaction | 1 | | 2.69 | 0.34 | | |
| | | | | | Sample size | n = 31 | | | |
| | | | | | Outliers removed | n = 2 | | | |
| Size of Dominant Individual | LM | average coral diameter | covariable | 1 | F-value | 6.33 | 0.02* | marginal | 0.207 |
| for *G. fuscoruber* | | group size | covariable | 1 | | 0.58 | 0.45 | | |
| fourth-root transformed | | location | factor | 2 | | 1.07 | 0.36 | | |
| (includes all coral species) | | coral diam x group size | interaction | 1 | | 0.21 | 0.65 | | |
| | | | | | Sample size | n = 31 | | | |
| | | | | | Outliers removed | n = 0 | | | |
| Coral Diameter | LM | size of dominant individual | covariable | 1 | F-value | 13.86 | 0.001* | marginal | 0.629 |
| for *G. fuscoruber* | | group size | covariable | 1 | | 23.94 | <0.0001* | | |
| log-transformed | | location | factor | 2 | | 1.69 | 0.21 | | |
| (includes all coral species) | | dominant size x group size | interaction | 1 | | 1.30 | 0.27 | | |
| | | | | | Sample size | n = 31 | | | |
| | | | | | Outliers removed | n = 0 | | | |

Supplementary Table 3. Statistical output of size ratio and sex dominance analyses. GLM = generalized linear model.

| Response Variable | Model | Predictor variable | Factor Type | df | Model Statistic | Test-value | p-value | R-squared statistic | Test value |
|---|---|---|---|---|---|---|---|---|---|
| Rankstep1 Size Ratios for | GLM | average coral diameter | covariable | 1 | χ2 (chi-squared) | 25.95 | 0.94 | | |
| single species colonies | quasibinom | group size | covariable | 1 | | 25.53 | 0.09 | | |
| without *G. brochus* | | location | fixed | 2 | | 25.34 | 0.52 | | |
| all locations | | species | fixed | 5 | | 23.08 | 0.15 | | |
| fourth power transformation | | location x species | interaction | 8 | | 21.56 | 0.24 | | |
| | | | | | Sample size | n = 164 | | | |
| | | | | | Outliers removed | n = 3 | | | |
| Rankstep1 Size Ratios for | GLM | average coral diameter | covariable | 1 | χ2 (chi-squared) | 26.17 | 0.21 | | |
| single species colonies | quasibinom | group size | covariable | 1 | | 25.98 | 0.25 | | |
| with *G. brochus* | | species | fixed | 6 | | 24.49 | 0.12 | | |
| all locations | | | | | Sample size | n = 175 | | | |
| fourth power transformation | | | | | Outliers removed | n = 5 | | | |
| Rankstep2 Size Ratios for | GLM | average coral diameter | covariable | 1 | χ2 (chi-squared) | 4.53 | 0.003* | | |
| single species colonies | quasibinom | group size | covariable | 1 | | 3.76 | 0.003* | | |
| all locations | | species | fixed | 3 | | 2.61 | 0.05 | | |
| | | | | | Sample size | n = 175 | | | |
| | | | | | Outliers removed | n = 0 | | | |
| Rankstep1 Size Ratios for | GLM | average coral diameter | covariable | 1 | χ2 (chi-squared) | 11.57 | 0.02* | | |
| single species colonies | quasibinom | group size | covariable | 1 | | 11.56 | 0.76 | | |
| between year at | | species | fixed | 4 | | 11.09 | 0.30 | | |
| LI and PNG only | | location | fixed | 1 | | 11.02 | 0.37 | | |
| fourth power transformation | | year | fixed | 1 | | 10.73 | 0.09 | | |
| | | species x location | interaction | 1 | | 10.41 | 0.07 | | |
| | | species x year | interaction | 4 | | 9.93 | 0.29 | | |
| | | location x year | interaction | 1 | | 9.93 | 0.81 | | |
| | | | | | Sample size | n = 114 | | | |
| | | | | | Outliers removed | n = 3 | | | |
| Rankstep1 Size Ratios for | GLM | average coral diameter | covariable | 1 | χ2 (chi-squared) | 16.79 | 0.001* | | |
| single species colonies | quasibinom | group size | covariable | 1 | | 16.34 | 0.06 | | |
| pre vs. post disturbance | | species | fixed | 3 | | 15.73 | 0.19 | | |
| LI only | | pre/post years | fixed | 2 | | 15.41 | 0.29 | | |
| fourth power transformation | | spp x pre/post years | interaction | 6 | | 14.32 | 0.20 | | |
| | | | | | Sample size | n = 124 | | | |
| | | | | | Outliers removed | n = 4 | | | |

Suppl. Tab. 3 (cont'd)

| Response Variable | Model | Predictor variable | Factor Type | df | Model Statistic | Test-value | p-value | R-squared statistic | Test value |
|---|---|---|---|---|---|---|---|---|---|
| Size Ratios among ranksteps | GLM | coral size | covariable | 1 | χ2 (chi-squared) | 40.83 | 0.002* | | |
| for mixed species colonies | quasibinom | group size | covariable | 1 | | 38.71 | 0.008* | | |
| fourth power transformation | | rankstep | fixed | 7 | | 35.36 | 0.14 | | |
| | | location | fixed | 2 | | 34.03 | 0.11 | | |
| | | coral size x rankstep | interaction | 7 | | 32.49 | 0.65 | | |
| | | group size x rankstep | interaction | 7 | | 30.13 | 0.35 | | |
| | | coral size x group size | interaction | 1 | | 29.92 | 0.41 | | |
| | | | | | Sample size | n = 119 | | | |
| | | | | | Outliers removed | n = 0 | | | |
| Size Ratios among ranksteps | GLM | coral size | covariable | 1 | χ2 (chi-squared) | 76.43 | 0.06 | | |
| for comparison of single vs. | quasibinom | group size | covariable | 1 | | 75.57 | 0.04* | | |
| mixed species colonies per | | rankstep | fixed | 6 | | 69.24 | < 0.0001* | | |
| species | | species | fixed | 6 | | 67.76 | 0.31 | | |
| fourth power transformation | | location | fixed | 2 | | 66.67 | 0.07 | | |
| | | mix/single colony | interaction | 1 | | 63.93 | < 0.0001* | | |
| | | location x mix/single | interaction | 2 | | 63.22 | 0.18 | | |
| | | rankstep x mix/single | interaction | 6 | | 61.36 | 0.18 | | |
| | | | | | Sample size | n = 307 | | | |
| | | | | | Outliers removed | n = 4 | | | |
| Size Ratios among ranksteps | GLM | coral size | covariable | 1 | χ2 (chi-squared) | 88.71 | 0.038* | | |
| for comparison of single vs. | quasibinom | group size | covariable | 1 | | 88.58 | 0.45 | | |
| mixed species colonies | | rankstep | fixed | 7 | | 77.97 | < 0.0001* | | |
| regardless of species | | location | fixed | 2 | | 77.06 | 0.13 | | |
| fourth power transformation | | mix/single spp colony | fixed | 1 | | 76.73 | 0.22 | | |
| | | location x mix/single spp | interaction | 7 | | 75.02 | 0.36 | | |
| | | coral size x rankstep | interaction | 7 | | 72.46 | 0.12 | | |
| | | group size x rankstep | interaction | 1 | | 72.29 | 0.36 | | |
| | | coral size x group size | interaction | 2 | | 71.39 | 0.13 | | |
| | | | | | Sample size | n = 338 | | | |
| | | | | | Outliers removed | n = 5 | | | |
| Sex Dominance divergence | type of test: 1-sample proportions test with continuity correction | | | | | | | | |
| from unity 1:1 | males: n = 62 AND females: n = 89 | | | 1 | χ2 (chi-squared) | 5.12 | 0.024* | | |
| Sex Dominance for | GLM | species | fixed | 4 | χ2 (chi-squared) | 4.88 | 0.30 | McFadden | 0.105 |
| single species colonies | binomial | year | fixed | 1 | | 13.48 | 0.0002* | Cox and Snell (ML) | 0.645 |
| at LI only: 2020 vs. 2021 | | species x year | interaction | 4 | | 4.97 | 0.29 | Nagelkerke | 0.645 |
| | | | | | Sample size | n = 162 | | | |

Supplementary Table 4. Statistical outputs of mixed species colony compositions. VGLM = multinomial logistic regression model.

| Response Variable | Model | Predictor variable | Factor Type | df | Model Statistic | Test-value | p-value | R-squared statistic | Test value |
|---|---|---|---|---|---|---|---|---|---|
| Mixed Species: Intermixed | VGLM | location | fixed | 2 | χ2 (chi-squared) | 0.19 | 0.91 | McFadden | 0.004 |
| Ranks Within Hierarchy? | | | | | | | | Cox and Snell (ML) | 0.005 |
| (only pre-disturbance data | | | | | | | | Nagelkerke | 0.007 |
| available):Yes or No | | | | | Sample size | n = 38 | | | |
| Mixed Species: Bigger Species | VGLM | location | fixed | 2 | χ2 (chi-squared) | 0.15 | 0.93 | McFadden | 0.004 |
| Is Rank 1? | | | | | | | | Cox and Snell (ML) | 0.004 |
| (only pre-disturbance data | | | | | | | | Nagelkerke | 0.006 |
| available):Yes or No | | | | | Sample size | n = 37 | | | |
| Mixed Species: Composition of | VGLM | location | fixed | 2 | χ2 (chi-squared) | 5.59 | 0.69 | McFadden | 0.110 |
| Colony | | | | | | | | Cox and Snell (ML) | 0.272 |
| only pre-disturbance | | | | | Sample size | n = 57 | | Nagelkerke | 0.288 |
| Mixed Species: Composition of | VGLM | pre vs. post disturbance | fixed | 2 | χ2 (chi-squared) | 2.89 | 0.58 | McFadden | 0.051 |
| Colony: OTI only | | | | | | | | Cox and Snell (ML) | 0.142 |
| pre vs post-disturbance | | | | | Sample size | n = 46 | | Nagelkerke | 0.149 |

Supplementary Table 5. *Gobiodon* species compositions in mixed species colonies.

| Mixed Species: Species Composition | Count |
|---|---|
| *G. axillaris & G. erythrospilus & G. fuscoruber & G. oculolineatus* | 1 |
| *G. axillaris & G. rivulatus* | 2 |
| *G. brochus & G. erythrospilus* | 2 |
| *G. brochus & G. rivulatus* | 5 |
| *G. c.f. fulvus & G. quinquestrigatus* | 1 |
| *G. citrinus & G. fuscoruber & G. okinawae* | 1 |
| *G. citrinus & G. okinawae* | 1 |
| *G. citrinus & G. sp.D* | 1 |
| *G. erythrospilus & G. fuscoruber* | 6 |
| *G. erythrospilus & G. fuscoruber & G. oculolineatus* | 1 |
| *G. erythrospilus & G. oculolineatus* | 1 |
| *G. erythrospilus & G. rivulatus* | 1 |
| *G. fuscoruber & G. oculolineatus* | 4 |
| *G. fuscoruber & G. oculolineatus & G. quinquestrigatus* | 1 |
| *G. fuscoruber & G. oculolineatus & G. rivulatus* | 1 |
| *G. fuscoruber & G. okinawae & G. quinquestrigatus* | 1 |
| *G. fuscoruber & G. quinquestrigatus* | 20 |
| *G. fuscoruber & G. quinquestrigatus & G. rivulatus* | 2 |
| *G. fuscoruber & G. rivulatus* | 9 |
| *G. histrio & G. oculolineatus* | 1 |
| *G. histrio & G. quinquestrigatus* | 2 |
| *G. histrio & G. rivulatus* | 3 |
| *G. oculolineatus & G. quinquestrigatus* | 1 |
| *G. oculolineatus & G. quinquestrigatus & G. rivulatus* | 2 |
| *G. oculolineatus & G. rivulatus* | 8 |
| *G. oculolineatus & G. sp.D* | 1 |
| *G. okinawae & G. quinquestrigatus* | 2 |
| *G. okinawae & G. sp.D* | 1 |
| *G. quinquestrigatus & G. rivulatus* | 3 |
| *G. sp.D & G. rivulatus* | 1 |

| Species | Count in Mixed Colony | Count in Single Species Colony |
|---|---|---|
| *G. axillaris* | 3 | 19 |
| *G. brochus* | 7 | 17 |
| *G. c.f. fulvus* | 1 | |
| *G. citrinus* | 3 | 3 |
| *G. erythrospilus* | 12 | 42 |
| *G. fuscoruber* | 47 | 45 |
| *G. histrio* | 6 | 44 |
| *G. oculolineatus* | 21 | 27 |
| *G. okinawae* | 6 | 10 |
| *G. quinquestrigatus* | 35 | 83 |
| *G. rivulatus* | 37 | 70 |
| *G. sp.D* | 4 | 5 |

| # Species in Mixed Colony | Count |
|---|---|
| Two species | 76 |
| Three species | 9 |
| Four species | 1 |
